# miR-155 exerts posttranscriptional control of autoimmune regulator (Aire) and tissue-restricted antigen genes in medullary thymic epithelial cells

**DOI:** 10.1101/2021.07.01.450769

**Authors:** Pedro Paranhos Tanaka, Ernna Hérida Oliveira, Mayara Cristina Vieira-Machado, Max Jordan Duarte, Amanda Freire Assis, Karina Fittipaldi Bombonato-Prado, Geraldo Aleixo Passos

## Abstract

**Background:** The autoimmune regulator (Aire) gene is critical for the appropriate establishment of central immune tolerance. As one of the main controllers of promiscuous gene expression in the thymus, Aire promotes the expression of thousands of downstream tissue-restricted antigen (TRA) genes, cell adhesion genes and transcription factor genes in medullary thymic epithelial cells (mTECs). Despite the increasing knowledge about the role of Aire as an upstream transcriptional controller, little is known about the mechanisms by which this gene could be regulated.

**Results:** Here, we assessed the posttranscriptional control of Aire by miRNAs. The *in silico* miRNA-mRNA interaction analysis predicted thermodynamically stable hybridization between the 3′ UTR of Aire mRNA and miR-155, which was confirmed to occur within the cellular milieu through a luciferase reporter assay. This finding enabled us to hypothesize that miR-155 might play a role as an intracellular posttranscriptional regulator of Aire mRNA. To test this hypothesis, we transfected a murine mTEC cell line with a miR-155 mimic *in vitro*, which reduced the mRNA and protein levels of Aire. Moreover, large-scale transcriptome analysis showed the modulation of 311 downstream mRNAs, which included 58 TRA mRNAs. Moreover, miR-155 mimic-transfected cells exhibited a decrease in their chemotaxis property compared with control thymocytes.

**Conclusion:** Overall, the results indicate that miR-155 may posttranscriptionally control Aire mRNA as well as a crucial process by which mTECs allow migration of thymocytes through chemotaxis.

## Background

Promiscuous gene expression (PGE) is a highly conserved process in mice and humans and is characterized by the ectopic expression of genes in the thymus and specifically in medullary thymic epithelial cells (mTECs), making mTECs the cell type known to express the highest number of genes to date [1-3]. PGE is a population-level process, that is, the total number of ectopic genes expressed is an estimate of the sum of the expression levels in each individual mTEC, and this process ensures self-representation and self-reactive T cell elimination through negative selection [4,5]. These processes guarantee that central tolerance is extended to virtually all tissues and organs in the body.

One of the main controllers of PGE in mTECs is the autoimmune regulator (Aire) gene, which encodes the AIRE protein that acts as a noncanonical transcription factor that promotes the expression of tissue-restricted antigens (TRAs) [6-9]. According to Takaba et al [10] 28.9% of the TRAs expressed in mTECs are under the control of Aire and are referred to as Aire-dependent genes. In addition, another 12.3% of TRAs are under the control of Aire and Forebrain Embryonic Zinc Finger-Like Protein 2 (Fezf2), and the other 58.8% are not under the control of Aire and are referred to as Aire-independent genes.

The Aire gene is expressed in only a few tissues, but it is mostly expressed in the thymus and, more specifically, in mTECs [11]. Additionally, only a subpopulation of mature mTECs with the CD80^hi^ MHC-II^hi^ phenotype express the Aire gene; these cells are referred to as mTEC^hi^ Aire^+^ cells.^1,5^ The AIRE protein acts as a transcriptional modulator of the chromatin of mTECs, promoting the expression of several Aire-dependent TRAs [2,3,11]. The importance of Aire in controlling the maturation and proliferation of mTECs, as well as its participation in the process of adhesion between mTECs and thymocytes during negative selection, have been shown [2,3,11-13].

Immature CD80^low^ MHC-II^low^ mTECs, which do not express Aire, express a smaller set of TRAs, while the mTEC^hi^ Aire^+^ cell population can express up to 62% of the protein-coding genes (mainly encoding TRAs) in mice [2,3,5].

In addition to TRAs, which are extremely important for testing the self-reactivity of thymocytes, studies have shown that several Aire-dependent genes have different functions in mTECs; these genes participate in processes such as the differentiation and maturation of mTECs,^14^ interactions of mTECs with thymocytes and dendritic cells [15,16], induction of thymocyte migration within the thymus, apoptosis of mTECs and even the processing of mRNAs (alternative splicing), thus contributing to the expression of TRA isoforms in mTECs [1-3,5,6,17].

Our group, and independently, the former group of Kyewski/Ucar, has contributed to showing that Aire is involved in the expression of miRNAs in mTECs [12,18]. Recently, it was shown that Aire transactivates the expression of HLA-G (19) and that it can regulate the length of the 3′
s untranslated regions (3′UTRs) of downstream mRNAs [20,21].

The AIRE protein is composed of five main subdomains: the CARD (caspase recruiter domain) domain, which acts in the homo-multimerization of the protein, since AIRE acts as a tetramer in the nucleus of mTECs; the NLS (nuclear localization signal) domain, which functions in the translocation of the protein to the nucleus; the SAND domain, which is involved in the protein-protein interactions that enable AIRE to act in conjunction with its partner proteins; and 2 PHD domains, namely, PHD-1 and PHD-1, separated by a proline-rich region (PRR), which act as a histone code reader and assists in chromatin decondensation [2,6,22].

AIRE acts as a transcriptional modulator because it does not recognize any particular motif in DNA to promote transcription. The AIRE protein interacts with the RNA Pol II molecule that has already started the transcription process. RNA Pol II is “anchored” to chromatin at specific T-box (TTATTA) and G-box (ATTGGTTA) sites in the promoter regions of Aire-dependent genes and is then released by AIRE, promoting the extension of the transcription of downstream genes [23].

AIRE cooperates with other proteins (called Aire partners) to promote the expression of Aire-dependent genes in mTECs. Sirtuin 1 (SIRT-1) promotes the deacetylation of the lysine residues of AIRE to promote its activation. Once activated, AIRE recognizes and binds to epigenetic markers involved in transcriptional repression, such as H3K4me0. Topoisomerase II (TOP2a) associates with AIRE and promotes double-strand DNA breaks. DNA PKs and other partners are activated by double-strand DNA breaks and promote the relaxation of chromatin. The protein complex of bromodomain-containing 4 (Brd4) and positive transcription elongation factor b (pTEFb) are associated with AIRE and play a role in releasing RNA Pol II from being anchored to chromatin. Then, CREB-binding protein (CBP) acetylates the lysine residues of AIRE, leading to their inactivation [2,6].

As introduced by Perniola [24], the expression of the Aire gene can be regulated by different mechanisms, including the lymphotoxin pathway (immunological manipulation) [25-27], viral infection [28,29], small-interfering RNA (siRNA) [16,30] or chemicals [31].

Aire, as stated above, is an important upstream transcriptional controller in mTECs, but little is known about the elements or mechanisms that control this gene. The structural characteristics of AIRE and its association with its partners control the molecular action of the AIRE protein in a posttranslational manner. Here, we ask which element(s) regulate the Aire gene posttranscriptionally, that is, after the transcription of Aire mRNA and when it is located in the cytoplasm. Considering the predicted significant thermodynamic stability of the interaction of the Aire mRNA 3′ UTR and miR-155, we hypothesized that Aire may be posttranscriptionally controlled in mTECs.

## Results

### Prediction of the miR-155-Aire 3′ UTR interaction and validation by luciferase reporter gene assay (LRGA)

The mmu-miR-155-5p (miR-155) has the following sequence: UUAAUGCUAAUUGUGAUAGGGGU. Analyses performed with the RNAhybrid algorithm predicted that the Aire mRNA 3′ UTR strongly hybridizes to this miRNA, exhibiting a minimal free energy (mfe) equal to -22 kcal/mol, which gives the structure thermodynamic stability (Figure 1).

**Figure 1.**
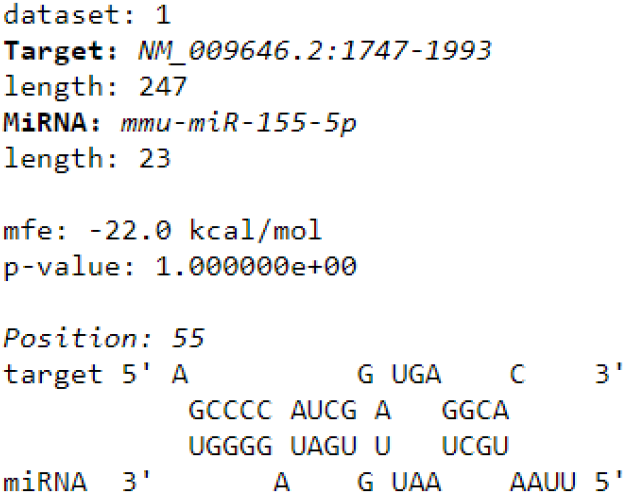
Prediction of posttranscriptional interaction between Aire mRNA 3′ UTR and miR-155 as evaluated by RNA-Hybrid program. The interaction resulted in a significant minimal free energy (-22 mfe) forming thus a stable hybrid structure.

The miRNA-mRNA interaction was then validated using the LRGA. The results showed that the luciferase expression decreased in cells cotransfected with the pMIR-Glo/Aire 3′ UTR vector and the miR-155 mimic. This result shows that there was an interaction between the miR-155 mimic and the 3′UTR of Aire mRNA (Figure 2).

**Figure 2.**
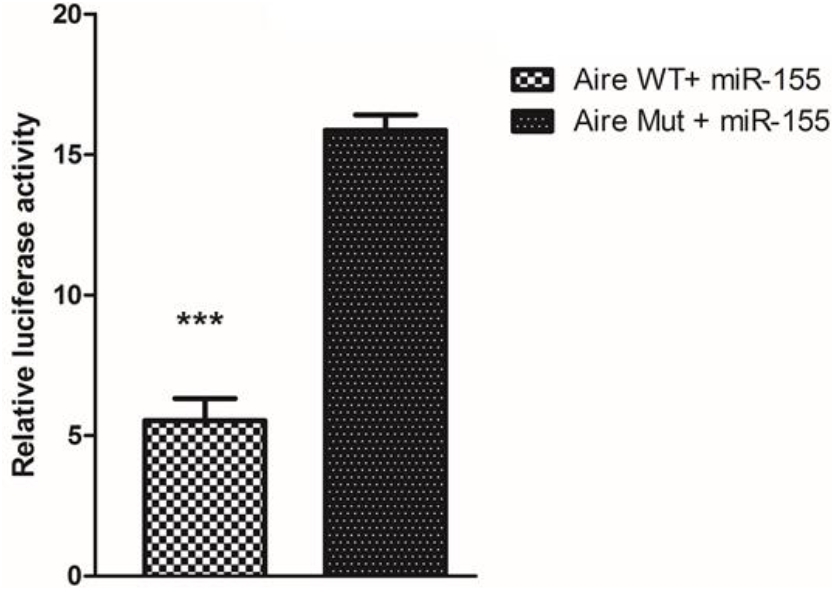
Posttranscriptional interaction between miR-155 with Aire 3′ UTR was assayed by luciferase reporter gene assay (LRGA). pMIR-Aire-wt-3′utr (wild type 3′UTR) or pMIR-Aire-mut-3′utr (mutant 3′UTR) luciferase vector constructs were transfected into human HEK-293T cells to demonstrate the possibility of occurrence of the miRNA-mRNA interactions within the cell milieu. The mutant 3′UTR was used to demonstrate the need of sequence specificity for the interaction to occur.

### Quantification of the intracellular levels of miR-155 and Aire mRNA

To assess the effect of miR-155 on Aire mRNA, mTEC 3.10 cells were transfected with the miR-155-5p mimic. RT-qPCR analysis of miR-155 was then performed to confirm the transfection efficiency and assess the intracellular levels of miR-155 at 12, 24, and 48 h posttransfection.

Compared with the control, transfection of the miR-155-5p mimic increased the intracellular level of miR-155, and peak expression was achieved at 12h posttransfection. Twenty-four hours posttransfection, the levels of miR-155 were notably decreased and were statistically comparable to the levels in the control cells. The results showed that 12 h posttransfection was the time point that was most appropriate for evaluating the effect of increasing the levels of miR-155 in mTEC 3.10 cells (Figure 3).

**Figure 3.**
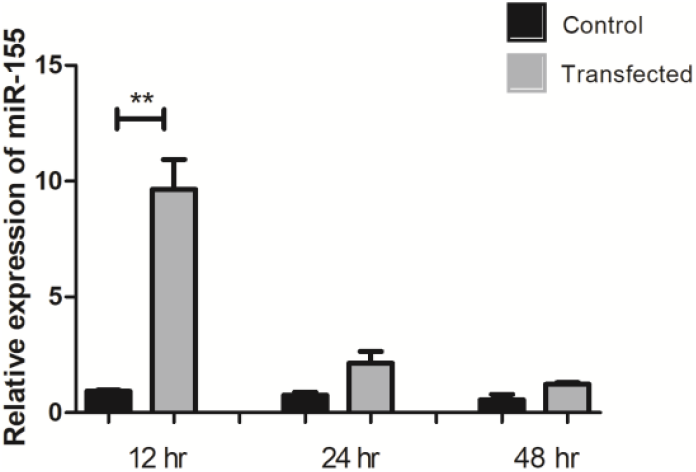
Relative expression of miR-155 in mTEC 3.10 cell line as detected by RT-qPCR. The mTEC cells were transfected (or not) with miR-155-5p mimic, causing significant increase in the levels of this miRNA 12 h after transfection. Difference between groups was analyzed by Student-t test, comparing data from control (not transfected) versus transfected cells (\*\**P <* 0.01).

Samples of total RNA (cDNA) from mTEC 3.10 cells transfected with the miR-155 mimic were used for RT-qPCR analysis of Aire mRNA. We again focused on the time points of 12, 24, and 48 h posttransfection. The results showed a significant reduction in Aire mRNA expression 12 h after the peak level of miR-155 expression was observed, that is, at 24 h posttransfection. Forty-eight hours after miR-155 transfection, the Aire mRNA levels were comparable to those in the control cells (Figure 4).

**Figure 4.**
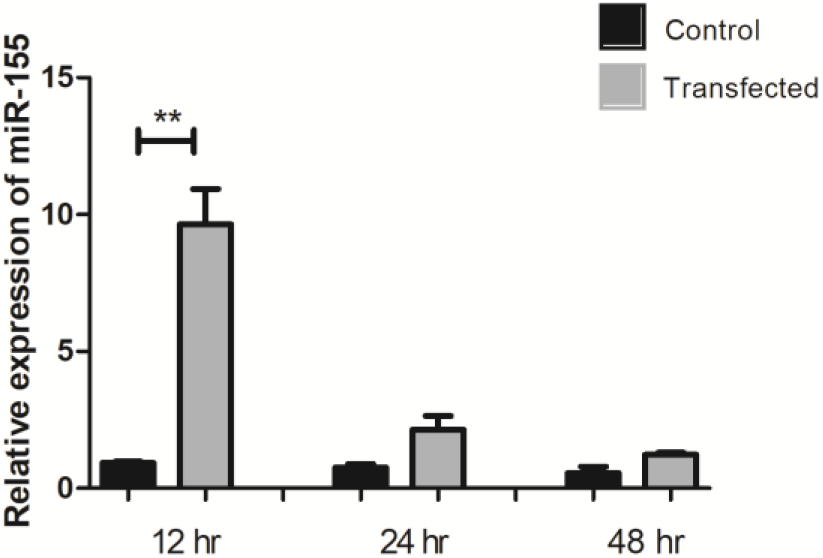
Relative expression of Aire mRNA in mTEC 3.10 cell line as detected by RT-qPCR. The mTEC cells were transfected (or not) with miR-155-5p mimic, causing significant decrease in the levels of this mRNA 24 h after transfection. Difference between groups was analyzed by Student-t test, comparing data from control (not transfected) versus transfected cells (\*\*\**P <* 0.001).

### AIRE protein expression analysis

#### Western blotting

The results obtained by RT-qPCR showed that the mRNA levels of Aire decreased 24 h after transfection with the miR-155-5p mimic. Thus, we decided to evaluate the protein levels of AIRE by western blot at this time point. AIRE protein expression in mTEC 3.10 cells was reduced (mean reduction of 37.3%) 24 h after transfection with the miR-155-5p mimic (Figure 5).

**Figure 5.**
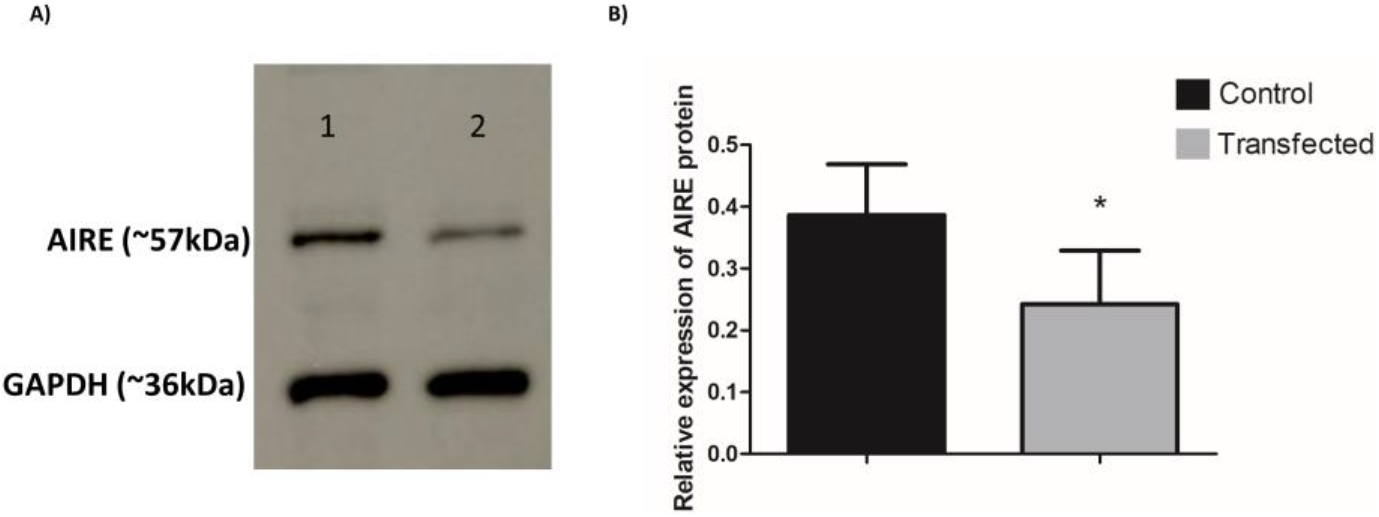
Expression of AIRE protein in mTEC 3.10 cell line as detected by western-blot. The mTEC cells were transfected (or not) with miR-155-5p mimic, causing significant decrease in the levels of this protein. (A) western-blot detection of AIRE after 24 h of transfection. The GAPDH protein was used as housekeeping and to normalize data. (B) Bar graph resulted from quantification of western-blot bands show the effect of miR-155-5p mimic transfection on AIRE expression. The data are presented as the means and standard error of mean (SEM) from three independent determinations. Difference between groups was analyzed by Student t-test, comparing control (not transfected) versus transfected cells (\*\**P <* 0.01).

#### Immunofluorescence

To evaluate the effect of miR-155-5p mimic transfection on AIRE protein levels in subcellular compartments, we used immunofluorescence. mTEC 3.10 cells were observed at 24 h posttransfection. Results show that the AIRE protein was located in the nucleus and visualized as red dots. Twenty-four hours posttransfection, there was a reduction of approximately 15% in the fluorescence signal intensity. The number of Aire^+^ mTEC 3.10 cells remained comparable to that of the control group. The results confirmed that transfection with the miR-155 mimic interfered with the AIRE protein expression level but not with the number of Aire^+^ mTECs (Figure 6).

**Figure 6.**
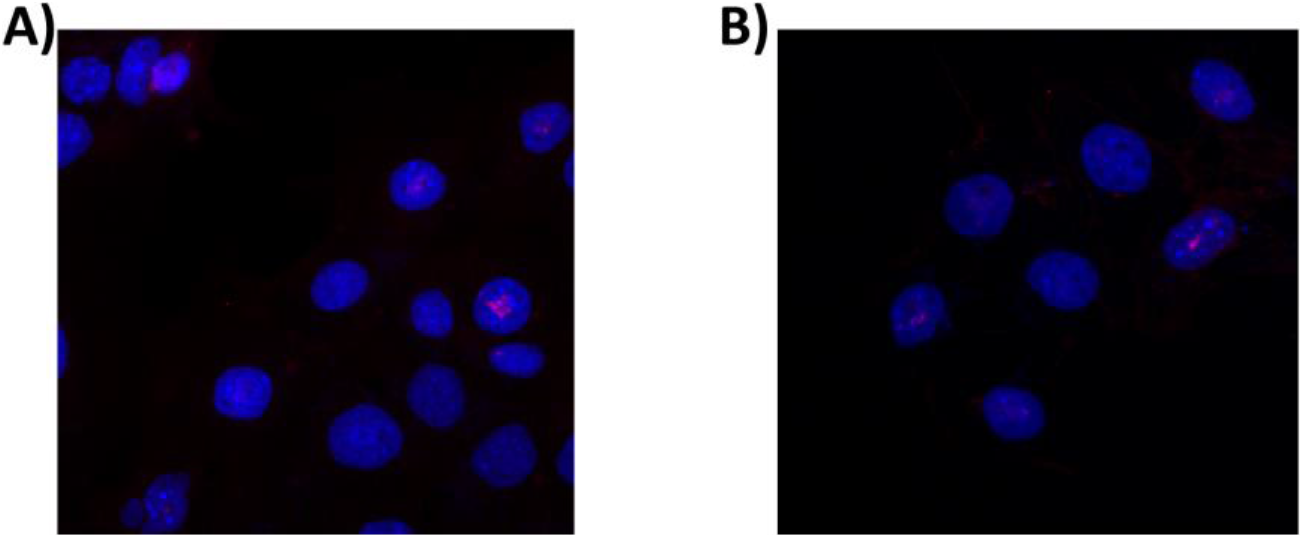
Expression of AIRE protein in mTEC 3.10 cell line as detected immunofluorescence. The mTEC cells were transfected (or not) with miR-155-5p mimic, causing significant decrease in the levels of this protein. (A) immunofluorescence detection of AIRE (red dots in the nuclei stained in blue) after 24 h of transfection. (B) Bar graph resulted from quantification of immunofluorescence signals show the effect of miR-155-5p mimic transfection on AIRE expression. The data are presented as the means and standard error of mean (SEM) from three independent determinations. Difference between groups was analyzed by Student t-test, comparing control (not transfected) versus transfected cells (\*\**P <* 0.01).

### Differentially expressed mRNAs in mTEC 3.10 cells transfected with the miR-155 mimic

The mTECs transfected with the miR-155-5p mimic exhibited the most significant effects on the mRNA and protein levels of Aire 24 h posttransfection. Therefore, these samples were selected for large-scale gene expression analysis (mRNA transcriptome) using the microarray technique and the limma linear model package for data analysis.

mRNAs were considered to be differentially expressed when they exhibited a fold change of ≥ 1.5. As a cutoff, an adjusted p-value ≤ 0.05 (Benjamini-Hochberg FDR) was used. The differentially expressed mRNAs were then visualized on a heatmap (Figure 7). In total, 311 mRNAs exhibited altered expressions; of these, 103 were upregulated, and 208 were downregulated. Of these 311 mRNAs, 249 encode previously described proteins. However, the other 62 corresponded to predicted mRNAs that have not been recorded in public databases.

**Figure 7.**
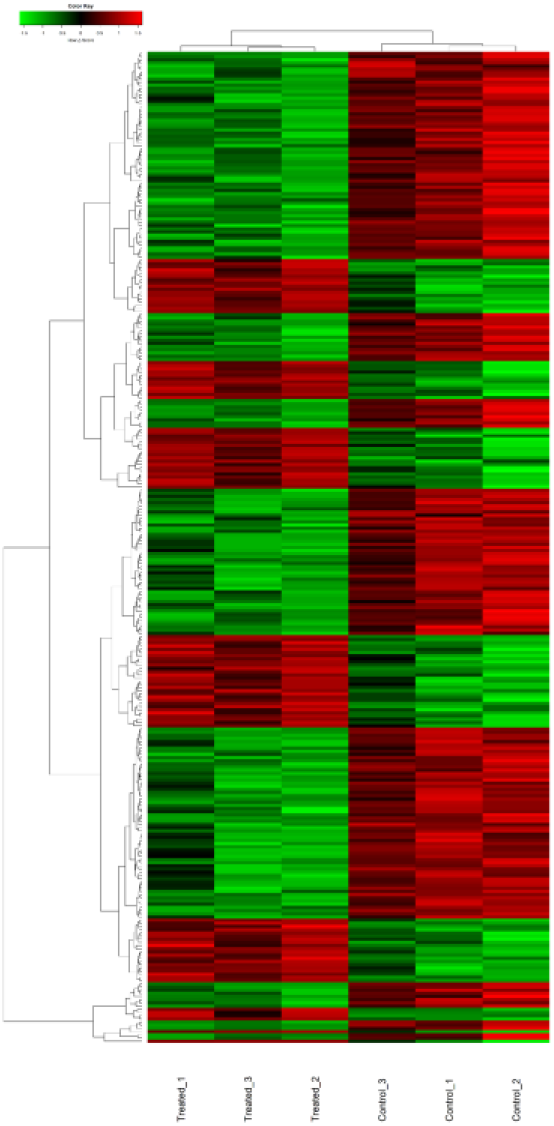
Transcriptome (mRNAs) expression profiles of mTEC 3.10 cell line transfected (or not) with miR-155-5p mimic. The mTEC cells total RNA samples were analyzed through microarray hybridizations, which allowed identify the differentially expressed mRNAs including those that encode tissue-restricted antigens (TRAs) or cell migration molecules. The dendrograms and heat-maps were obtained using the cluster and tree view algorithm considering 1.5-fold-change and 0.05 false discovery rate. Heat-map legend: red = upregulation, green = downregulation, black unmodulation (Pearson’ s correlation metrics).

Of the 249 altered and annotated mRNAs, 58 (23.3%) were considered to encode tissue-restricted antigens (TRAs). Compared to the average expression levels, the expression of these mRNAs was increased in a maximum of five tissues [1]. As miRNAs have several mRNA targets, in this work, we evaluated which of the mRNAs with altered expression could be targeted by miR-155 according to MirWalk predictions (Table 1).

**Table 1.** The mRNAs predicted by MirWalk as target of miR-155.

|  |  |
| --- | --- |
| 2010107E04Rik | Hapln3 |
| 2310079G19Rik | Kpna4 |
| 4930519N16Rik | Mccc2 |
| Ampd3 | Ndn |
| Bach1 | Prkci |
| Cycs | Rreb1 |
| Dbn1 | Snrpn |
| Gbe1 |  |

### Functional enrichment of differentially expressed mRNAs

To identify the effect of miR-155 on mTEC 3.10 cells, *in silico* predictions of the functions of the differentially expressed mRNAs was performed using the DAVID genome database platform (https://david.ncifcrf.gov/). Initially, the mRNAs were separated into two groups, namely, the upregulated and downregulated groups, and they were grouped based on their functions in biological processes, cellular components, or molecular function. Categories that included at least three mRNAs and a score < 0.05 with Benjamini-Hochberg correction were further considered (Figure 8).

**Figure 8.**
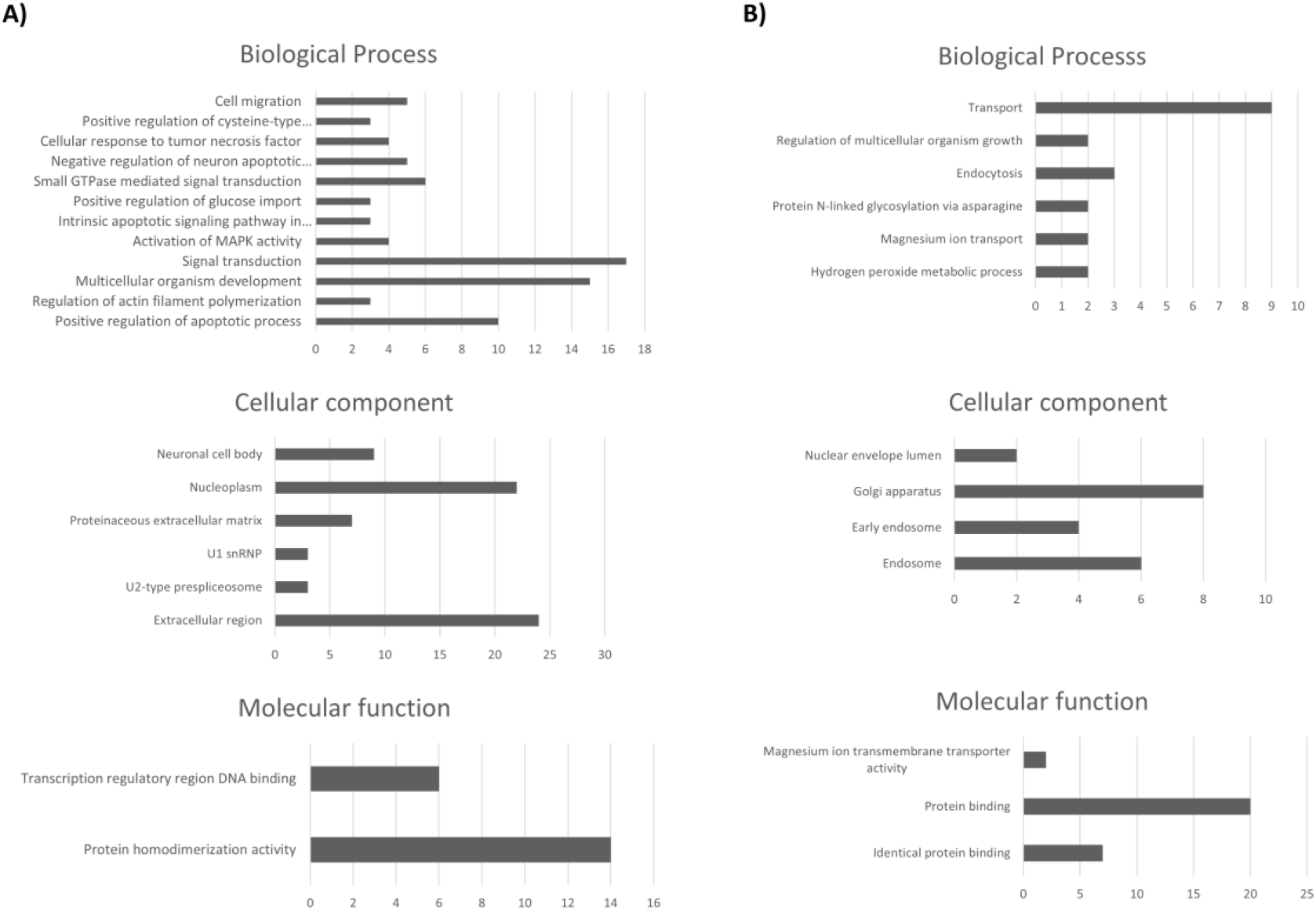
Functional annotation for modulated (up or downregulated) mRNAs in mTEC 3.10 cells. The functional categories were identified by using DAVID genome database platform according to GO processes: biological processes (BP), cellular components (CC) and molecular function (MF). The rank is based on the enrichment score, which represents mean *p*-value. Only those mRNA-groups yielding < 0.05 Benjamini corrected *p*-value and containing at least three mRNAs are considered to be significant.

### Thymocyte migration

To evaluate the effect of the miR-155-5p mimic on the chemotactic properties of mTEC 3.10 cells, we performed a thymocyte migration assay since five of the downregulated mRNAs (Erbb4, Efna1, Prkcl, Prss37, and Matk) are related to cell migration. Conditioned culture media from the control or miR-155-5p mimic-transfected mTEC cell cultures (24 h posttransfection) were used. It was possible to observe a reduction of approximately 34.7% in the number of thymocytes that migrated through the inserts when cultured with conditioned culture medium from transfected cells compared to conditioned culture medium from control cells (Figure 9).

**Figure 9.**
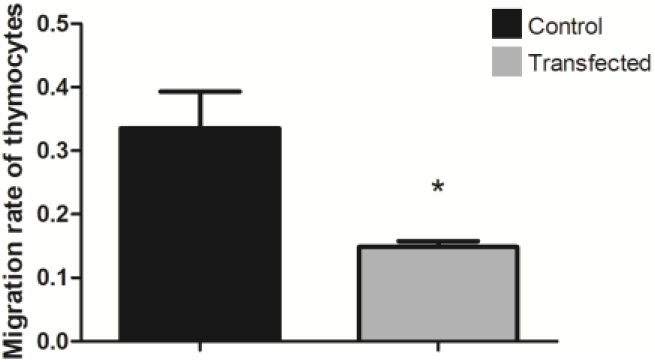
Decrease of thymocyte migration towards mTEC 3.10 cells (chemotaxis) following miR-155-5p mimic transfection. Differences in the numbers of migrated thymocytes through porous membrane in conditioned medium from transfected or not transfected cells were quantified, and data are shown in bar graph form. The migration degree is plotted as migration index, in which values correspond to mean ± s.d. Data are representative of three independent experiments and were significantly different between control cells versus cells transfected with miR-155-5p mimic. (\*\**P <* 0.01, Student t-test).

## Discussion

In this study, we investigated the posttranscriptional control of the autoimmune regulator (Aire) gene using an *in vitro* model system with a murine medullary thymic epithelial cell line. The control of the promiscuous gene expression (PGE) of autoantigens in mTECs is mainly performed by the Aire transcriptional regulator, along with the Fezf2 transcription factor, but to date, little is known about the mechanisms by which this gene is controlled [2,4,32].

Two important axes that control the expression of eukaryotic genes, which are controlled by transcription factors at the chromatin level [33] or posttranscriptionally by miRNAs in the cytoplasm [34-36], might act in this case. miRNAs finely control the expression of proteins by interacting with their respective mRNAs.

Here, we explored mechanisms of posttranscriptional control. We hypothesize that Aire is regulated by miR-155 in mTECs. This hypothesis was generated after we performed *in silico* prediction of miRNA-Aire mRNA interactions and observed a strong hybridization of miR-155 with the 3′ UTR of the mRNA that encodes the AIRE protein (Supplemental Figure 1).

This prediction was validated by a luciferase assay, making it possible to observe the miRNA-mRNA interaction in the intracellular milieu of human HEK-293T cells. However, we were interested in validating the miR-155-Aire mRNA interaction in the mTEC cell milieu.

We used the murine Aire^+^ mTEC 3.10 cell line [37], which was transfected with the miR-155-5p mimic. In the short term, this transfection led to a reduction in the protein expression of AIRE, as detected by western blotting. The AIRE protein levels returned to normal level, comparable to the levels in the control cells, 48 h after the transfection of the miR-155-5p mimic. The analyses showed that the dose and time course of the miR-155-5p mimic were sufficient to affect the mRNA and protein levels of Aire, which suggests possible consequences for the transcriptome of these cells due to the importance of Aire in PGE in mTECs.

Different time points after transfection of the miR-155-5p mimic, as well as different concentrations of the miR-155-5p mimic, were tested. Twelve hours of transfection with 2 nM miR-155-5p mimic proved to be the best conditions that achieved the highest concentration of intracellular miR-155-5p. At 24 h after the transfection of the miR-155-5p mimic, there was a drastic reduction in the level of Aire transcriptional expression, suggesting degradation of Aire mRNA by the RISC complex.

The levels of miR-155 returned to normal, that is, to levels comparable to those observed in control cultures, 24 h after transfection. Regarding the mRNA levels of Aire, we observed that 48 h after transfection, there was no longer a significant difference in its expression between the control and transfected cultures. This defined a “window” for making posttransfection observations: 12 h for miR-155 or 24 h for Aire mRNA. This window may be a result of the model system, which is analogous to small interfering RNA (siRNA), whose target inhibition is partial and transient [35,38]. However, the levels of overexpression of miR-155 and inhibition of Aire mRNA achieved in this model system were sufficient to answer the central question of the work.

Immunofluorescence showed no change in the number of Aire^+^ mTECs after transfection with the miR-155-5p mimic, but a reduction in the intensity of AIRE protein expression in each cell was observed. This result is consistent with the interpretation of an inhibitory effect of miR-155 on the translation of Aire mRNA rather than an elimination of Aire^+^ cells.

After observing the reduction in the mRNA and protein expression of Aire after the transfection of the miR-155-5p mimic, we proceeded to the analysis of the gene expression of mTEC 3.10 cells on a large scale, considering the impact of Aire and miR155-5p itself on the transcriptome of these cells.

Total RNA extracted from mTECs 24 h after transfection or from control cells was used to analyze gene expression by the microarray method.

We observed the modulation of 311 downstream mRNAs, of which, 103 were induced and 208 were repressed. Of the 311 modulated mRNAs, 249 were annotated in the DAVID genome database platform. Of these, 58 (23.3%) encode TRAs, which is consistent with Sansom Sans et al [1], which showed that approximately 25% of Aire-dependent genes encode TRAs.

Interestingly, of the 58 TRA mRNAs, ten were overexpressed in mTEC 3.10 cells transfected with the miR-155-5-mimic. These mRNAs must represent a portion of genes that, instead of being induced, are suppressed by Aire at the transcriptional level [1,5].

Considering that miRNAs, because they are small single-stranded RNAs, can have several targets, we evaluated which of the differentially expressed mRNAs might be targets of miR-155 are as predicted by the MirWalk algorithm.

Fifteen mRNAs were identified, of which three were predicted but none annotated, and another 12 were predicted and annotated; these mRNAs included Ampd3, Bach1, Cycs, Dbn1, Gbe1, Hapln3, Kpna4, Mccc2, Ndn, Prkci, Rreb1, and Snrpn. Of these, three (Bach1, Cycs, and Gbe1) were induced, while the others were repressed, indicating a direct action of miR155 on the 3′ UTRs of these nine mRNAs.

The results showed that the reduction in Aire expression caused by transfection with the miR-155-5p mimic was sufficient to cause the modulation of TRA mRNAs, which are essential for self-representation and negative selection in the thymus [2,39].

Many studies have shown that the importance of Aire in mTECs extends beyond the control of TRA expression during PGE [1,2,5,6,16,17]. The functional enrichment of the differentially expressed genes allowed us to identify mRNAs related to cell migration.

Pezzi et al [16]. showed that Aire controls the expression of mRNAs that encode proteins associated with cell adhesion (mTEC-thymocyte interaction), a process that is essential for negative selection. Among the downregulated mRNAs, we identified Cldn6 and Hapln3, which are both related to the cell adhesion process.

St-Pierre et al [5] demonstrated that the deletion of Aire in cTECs or in mTECs ^low^, that is, cells that do not express Aire, does not alter the complexity of splicing in these cells. Thus, AIRE can also participate in mRNA processing and contribute to isoform diversity of both Aire-dependent and Aire-independent TRA genes [17,40]. In this study, we identified mRNAs that encode proteins that participate in spliceosome assembly, such as SNRPN, PEPF39, SF3A2, and SNRPA.

Another possible role of Aire is in inducing the apoptotic process in mTECs. Although this function has not yet been fully confirmed, the induction of apoptosis by Aire would explain the rapid turnover of Aire^+^ mTECs and suggests that after the apoptotic process, these cells would be phagocytosed and that TRAs would be presented by antigen-presenting cells present in the thymic medulla, such as macrophages or dendritic cells, which assist in the negative selection process [2,6,41,42].

It was also possible to identify several inhibited mRNAs that act in the positive regulation of the apoptotic process, namely, Erbb4, Ddx19a, Muc20, Picard, Rhob, Pawr, Trp73, Cdk4, Sirt1, and Lats1.

Among the downregulated mRNAs, six were identified as being related to MAPK (mitogen-activated protein kinase) activation, such as Erbb4, Rreb1, Dmtf1, Trp73, Tbl1x, and Brca1. MAPKs regulate a diversity of biological processes, such as cell differentiation and apoptosis.

The mRNAs encoding ADAMTSL5 and ARSI proteins were also inhibited, and these mRNAs participate in protein metabolism and macromolecule degradation, respectively, which are crucial processes in mTECs since these cells express a vast array of ectopic proteins.

We also observed the repression of mRNAs related to the cell migration process, including Erbb4, Efna1, Prkcl, Prss37, and Matk. Laan et al [43]. showed that Aire increases the expression of chemokine ligands involved in the migration of thymocytes, such as CCR7 and CCR4, strongly suggesting a role for Aire in the chemotactic properties of mTECs and possibly influencing, even if indirectly, the migration of thymocytes to the thymus and within the thymus [13].

This prompted us to analyze the biological (functional) effect caused by transfection with the miR-155-5p mimic. The rationale was that if the transfection of miR-155-5p mimic causes a reduction in the expression of mRNAs that encode migration-related proteins, the chemotactic properties of the treated mTECs would be altered.

To test this hypothesis, we performed a thymocyte migration assay. The assay was carried out using inserts from the transwell culture plates that allow thymocytes to pass towards conditioned culture media (media from cultures of control mTECs or miR-155-5p mimic transfected mTECs). The transwell migration process was stopped after three hours to fix and stain the thymocytes in the inserts during the migration process and thus evaluate the migration rate.

Transfection of mTECs with the miR-155-5p mimic effectively reduced the mRNA and protein expression of Aire. The consequence of this was significant for the transfected cells, as we found hundreds of downstream mRNAs that were also reduced due to the reduction in Aire. Of the 311 reduced downstream mRNAs, only 15 are predicted targets of miR-155.

The results showed a significant reduction in thymocyte migration in the conditioned medium compared with cells transfected with the miR-155-5p mimic.

## Conclusions

The results showed that Aire might be regulated at the posttranscriptional level by miR-155. Data confirms, at the biological level and using an *in vitro* model system, that the decrease in the expression of Aire and consequently in the expression of mRNAs that encode chemotactic proteins might have functional consequences regarding the chemotactic property of mTECs.

## Material and Methods

### In silico predictions of miRNA-mRNA interactions

For the analysis of possible miRNAs that interact with Aire mRNA, a search was performed in public databases to predict miRNA-mRNA interactions using the MirWalk platform, which is available online at (http://zmf.umm.uni-heidelberg.de/apps/zmf/mirwalk2/miRretsys-self.html). The interaction of miR-155 with the 3′ UTR of Aire mRNA was predicted using the PITA algorithm, which is available at https://genie.weizmann.ac.il/pubs/mir07/mir07_prediction.html.

The thermodynamic stability of the molecular interaction was then validated using the RNAhybrid algorithm (https://bibiserv2.cebitec.uni-bielefeld.de/rnahybrid), which analyzes the secondary structure of the interaction and determines the most favorable hybridization site between a determined miRNA and the exact sequence of the 3′ UTR of its target mRNA. We only selected interactions with Gibbs free energy ≤ -20 kcal/mol, which are the most stable.

The algorithms of the MirWalk platform were used to predict other mRNAs, in addition to Aire mRNA, which could be targeted by miR-155.

### Validation of miRNA-mRNA interactions through luciferase reporter gene assay (LRGA)

In this study, we used an oligonucleotide encompassing the 3′ UTR of Aire mRNA synthesized through GBlocks® technology by Integrated DNA Technologies (IDT, Coralville, IA, USA). The oligonucleotide was cloned into the polycloning site of the pmirGLO vector (Promega Corporation) between the XhoI/XbaI restriction sites, resulting in the miRNA target region assuming the correct 5′ to 3′ orientation immediately downstream of the luciferase gene.

For the selected target, we introduced a point mutation into the 7-nucleotide seed-binding sequence. The constructs, named “pMIR-Aire-wt-3′ utr” to indicate the inclusion of the wild-type sequence and “pMIR-Aire-mut-3′utr” to indicate the inclusion of the mutant sequence, were selected by colony polymerase chain reaction (PCR) using a pair of primers flanking the vector polycloning site. We used *Escherichia coli* DH5α for cloning.

For the LRGA, 0.2 μg of each pmirGLO construct was transfected into human HEK-293T cells [44] (6 × 10^4^ cells/well) with 1.6 pmol of miR-155 or scrambled control miRNA (Thermo Scientific Dharmacon, Waltham, MA) in a 96-well plate. Transfections were performed using the Attractene Transfection Reagent (Qiagen, Hilden, Germany) according to the manufacturer’s instructions. The transfected cells were incubated at 37°C in a 5% CO_2_ incubator; 24 h after transfection, the cells were lysed in Passive Lysis Buffer. The firefly and Renilla luciferase activities were measured in a Synergy 2 luminometer (BioTek Instruments, Inc., Winooski, VT) using the Dual-Luciferase Reporter System (Promega Corporation) according to the manufacturer’s instructions.

The LRGA results are presented as the standard error of the mean (SEM). The differences were evaluated by one-way ANOVA followed by Student’s *t*-test (two groups). *P* < 0.05 was considered to be statistically significant.

Our laboratory has national biosafety approval (National Technical Committee for Biosafety, Brasilia, Brazil, CTNBio Permit No. 0040/98).

### mTEC 3.10 cell line, miR-155-5p mimic transfection and total RNA extraction

We used the murine mTEC 3.10 (EpCAM^+^, Ly51^−^, UEA-1^+^) cell line, which expresses wild-type (WT) *Aire*, as previously described [16,45,46].

For the transfection of mTEC 3.10 cells with the miR-155-5p mimic, we used an oligonucleotide synthesized by IDT whose sequence was 5′UUAAUGCUAAUUGUGAUAGGGG3′. The cells were transfected using Lipofectin Reagent from Gibco-BRL Life Technologies (Van Allen Way Carlsbad, CA, USA) following the manufacturer’s protocol. The mTEC 3.10 cells were grown in RPMI medium and 10% inactivated fetal bovine serum (FBS) at 37°C in a 5% CO_2_ atmosphere in T25 polystyrene flasks until they reached 50% confluence.

The miR-155-5p mimic-lipofectamine mixture was prepared by mixing 1 μL of 10 μM miR-155 mimic with 500 μL of Opti-MEM® I Reduced Serum Medium Gibco (Paisley, Scotland, UK). Next, 10 μl of Lipofectin Reagent was added. This mixture was vigorously stirred for 10 seconds and then incubated for 10 minutes at room temperature.

The miR-155-5p mimic-lipofectamine complex was added to the culture flasks in a total volume of 5 ml per bottle, resulting in a concentration of 2 nM miR155 mimic. For the control, mTEC 3.10 cells were cultured under similar conditions, except that the miR-155 mimic was added alone. The cells were cultured for different periods (12, 24, and 48 h posttransfection). Independent experiments and controls were repeated three times.

Total RNA was extracted from control or transfected cells using the mirVana miRNA Isolation Kit (Ambion, Grand Island, NY, USA) according to the manufacturer’ s instructions. The integrity of the RNA samples was evaluated through microfluidic gel electrophoresis in RNA 6000 nanochips and in an Agilent 2100 Bioanalyzer® device (Agilent Technologies, Santa Clara, CA, USA). We used only RNA preparations with an RNA integrity number (RIN) ≥ 7.5 that were free of phenol (A_260_/A_220_ approx. 0.7) and free of protein (A_260_/A_280_ approx. 2.0).

### Reverse transcription for cDNA synthesis

mRNA reverse transcription reactions were performed with components of the Applied Biosystems™ (Foster City, CA, USA) High Capacity cDNA Reverse Transcription Kit following the manufacturer’s protocol. For these reactions, 500 ng of total RNA was used. For the reverse transcription of miR-155, the TaqMan MicroRNA Reverse Transcription Kit (Foster City, CA) was used following the manufacturer’s protocol. For these reactions, 50 ng of total RNA was used.

### Real-time quantitative PCR (RT-qPCR)

The expression levels of Aire mRNA and miR-155 were assessed by reverse transcription quantitative real-time polymerase chain reaction (RT-qPCR). For these reactions, TaqMan Gene Expression Assay probes specific for Aire, Gapdh (endogenous control for mRNA), miRNA-155, and snoRNA-202 (endogenous control for miRNA) were used. The TaqMan Fast Advanced Master Mix kit (Applied Biosystems™) was used following the manufacturer’s protocol.

The expression level of miR-155 was normalized to that of snoRNA202, which is commonly used as an endogenous control for miRNA expression analysis.

The expression levels of the Aire gene were normalized to those of the constitutive gene Gapdh (ENSMUSG00000057666), which is commonly used as a reference.

The relative quantification of gene expression was performed using the 2-ΔΔCT method [47]. For the statistical analysis of the data, Student’s t-test and one-way ANOVA were performed using GraphPad Prism statistical software (www.scolary.com/tools/graphpad-prism) according to the manufacturer’s instructions.

### Extraction and quantification of total proteins from mTEC 3.10 cells

Approximately 5×10^5^ cells were seeded in T75 culture polystyrene flasks containing 10 mL of RPMI 1640 medium and 10% fetal bovine serum (FBS). The cells were transfected with the miR-155-5p mimic and cultured for 24, 36, or 48 h. The cells were detached from the culture flasks by the conventional trypsin treatment. The cells were washed three times with 1x PBS and centrifuged at 1000 x g for 5 minutes. Then, 300 μL of modified RIPA buffer [50mM TrisCl pH 7.4, 150mM NaCl, 1mM EDTA, 0.1% SDS, 0.1% protease inhibitor cocktail (Sigma-Aldrich, Darmstadt, Hesse, Germany)], and the cells were lysed using a Polytron homogenizer. The homogenate was then centrifuged at 10,000 x g for 20 minutes at 4°C. The total protein content of the supernatant was determined by using conventional Bradford reagent (Bio Rad, Hercules, CA, USA).

### Western blot analysis of the AIRE protein

The electrophoresis of total proteins in denaturing polyacrylamide gels (SDS-PAGE) was performed using the discontinuous buffer system described by Laemmli and Favre [48]. Polyacrylamide mini gels (10 × 8 cm; 0.075 cm thick) were used. The concentration of the stacking gel was 4%, and the concentration of the separation gel was 10%. The protein samples were prepared for electrophoresis at a ratio of 3:1 (vol/vol) of sample to 4x concentrated sample buffer (240 mM Tris-HCl pH 6.8; 8% SDS; 40% glycerol; 20% 2-mercaptoethanol and bromophenol blue). The samples were heated for 5 minutes at 95°C in a heat block. The electrode buffer contained 25 mM Tris-HCl, pH 8.3; 192 mM glycine; and 0.1% SDS. The protein molecular weight standard Precision Plus Protein ™ Dual Color Standards (BioRad, Hercules, CA, USA) was used.

The protein bands, once separated, were transferred from the gel to a PVDF membrane (BioRad, Hercules, CA, USA) using 25 mM Tris-HCl, 190 mM glycine, 20% methanol, and 1% SDS as transfer buffer at 80 volts for two hours. For this step, we used the BioRad Mini Trans-Blot® Cell device (Hercules, CA, USA). After the transfer, the membrane was stained with 0.2% Ponceau S in 3% trichloroacetic acid for 5 minutes to evaluate the transfer efficiency. Then, the membrane was destained with deionized water. The membrane was blocked with skim milk (MOLICO) diluted to 5% in TBS-Tween (50 mM Tris-HCl, pH 8.0; 150 mM NaCl; 0.1% Tween-20) for one hour at room temperature. After blocking, the membrane was incubated with the polyclonal anti-Aire D17 (goat antibody) primary antibody from Santa Cruz Biotechnology®, Inc. (Dallas, Texas, USA), which was diluted in TBS-T (1:500), for 18 h, followed by three washes in TBS-T for 5 minutes each.

The membrane was incubated with the anti-goat IgG secondary antibody conjugated to peroxidase (horseradish-peroxidase - HRP), which was diluted 1:10,000 in TBS-T, for 1 hour. After incubation, the membrane was washed in TBS-T, as before. The development was performed in an ImageQuant™ LAS 500 Ge Life Sciences apparatus (Piscataway, NJ, USA) using the appropriate substrate of peroxidase (Luminata™Forte, Merck Millipore). The reaction was conducted in the dark, and we exposed the membrane for different time intervals. The conversion of the intensity of the protein bands in the images into numerical values was performed with the aid of the ImageJ 1.49 program (http://imagej.nih.gov/ij/index.html). The GAPDH (glyceraldehyde 3-phosphate dehydrogenase) protein was used as an internal reference in the analyses. Thus, the membrane was also incubated with the polyclonal (rabbit) anti-GAPDH primary antibody from Cell Signaling Technology (Beverly, MA, USA). In that case, a secondary anti-rabbit IgG antibody conjugated to peroxidase was used. The immunostaining procedure was the same as that used for Aire.

### Immunolocalization of the AIRE protein

To assess the intracellular localization of the AIRE protein, we used the immunofluorescence of mTEC 3.10 cells (control or transfected with the miR-155 mimic). For this experiment, we used the anti-AIRE D17 primary antibody from Santa Cruz Biotechnology® (Dallas, Texas, USA). For this assay, 3×10^4^ mTECs (3.10) were plated on 13-mm^2^ glass coverslips placed in a 24-well plate. After transfection with the miR-155-5p mimic, the culture medium was removed from the wells, and the coverslips were washed five times with PBS.

The cells were fixed on the coverslips with 300 μL of 4% paraformaldehyde in PBS for 15 minutes. The cells were then subjected to three washes with PBS for five minutes each and then permeabilized with PBS containing 0.5% Triton X-100 for five minutes.

After three more washes with PBS (5 minutes each), the cells were blocked for one hour with 2% BSA in PBS. A diluted (1:50 in PBS containing 1% BSA) primary antibody (anti-AIRE D17) was added, followed by incubation at room temperature for one hour. The cells were washed three times with PBS (10 minutes each). Then, the Novex TM mouse anti-goat IgG (H + L) rhodamine red-conjugated secondary antibody (Life Technologies Corporation, Carlsbad, CA, USA), which was diluted 1:500 in PBS containing 1% BSA, was added and incubated for one hour. The coverslips were washed with PBS (3 × 10 minutes).

To visualize the cytoplasmic region, the actin filaments were labelled with phalloidin conjugated to AlexaFluor® 488 (Life Technologies™; Eugene, Oregon, USA) according to the manufacturer’s instructions. The coverslips were mounted with ProlongGold® Antifade Mountant (Life TechnologiesTM - Eugene, Oregon, USA) containing DAPI (to visualize the nuclei). The coverslips were visualized and photographed using an ApoTome Zeiss fluorescence microscope (Oberkochen, Germany).

### Microarray hybridizations and data analysis

This study followed a protocol that was previously established in our laboratory for microarray hybridizations [30]. Briefly, total RNA preparations were used to synthesize dscDNAs from mRNAs, and then, cyanine 3 (Cy3)-CTP-labeled complementary amplified RNAs (cRNAs) were synthesized. For these reactions, we used the Agilent Linear Amplification kit (Agilent Technologies) according to the instructions of the manufacturer. The (Cy3)-cRNA samples were hybridized to Agilent mouse 4 × 44 K-format oligonucleotide microarrays (Agilent Technologies). The slides were washed according to the manufacturer’ s instructions and scanned with an Agilent DNA microarray scanner.

The hybridization signals from the scanned microarray slides were extracted using Agilent Feature Extraction software, version 10.7.1.1. The expression profiles of independent samples of the control or miR-155-5p mimic-transfected mTEC 3.10 cells were analyzed by comparing the microarray hybridization signals of the respective samples. The numerical and quantitative microarray data were normalized to the 75^th^ percentile and analyzed in the R statistical environment (version 3.3.1) (https://www.r-project.org). In the data preprocessing phase, we used the arrayQualityMetrics [49] and Agi4×44PreProcess [50] tools, which have algorithms that allow the qualitative evaluation of arrays in addition to the correction of the background and normalization of the data.

For the analysis of differentially expressed mRNAs, we used the functions of the Limma package [51], which applies a linear model to the gene-wise statistical analysis. For an analysis of multiple tests, empirical Bayes and the Benjamini-Hochberg correction were applied.

In this work, we considered a p-value ≥ 0.05 with correction by FDR (Benjamini-Hochberg) and fold-change ≥ 1.5 to indicate differentially expressed mRNAs.

The differentially expressed transcripts were then subjected to hierarchical clustering, and heat maps were constructed to evaluate the expression pattern of the mRNAs. The Euclidean distance and the complete linkage method were used to group the samples and the RNAs.

### Thymocyte migration assay

For the separation of thymocytes, female C57BL/6 mice aged 5-6 weeks old were killed in a CO_2_ chamber, and the thymuses were quickly removed by thoracic surgery. The organs were fragmented with the aid of two surgical forceps in RPMI 1640 culture medium in a Petri dish. The thymocytes were separated through a 10 μm pore size nylon membrane (Sefar Inc. Depew, NY, USA). The thymocyte suspension was centrifuged for 5 minutes at 1000 × g, and the cells were washed twice with PBS and then resuspended in RPMI 1640 medium.

For the thymocyte migration assay, we used a protocol previously published [52] with modifications as follow. Transwell chambers and 6.5 mm inserts with a 0.5 μm in diameter polycarbonate membrane (Corning Inc. Corning, NY, USA) were used. The inserts were placed in 24-well plates and incubated in PBS for 1 hour at 37°C and then blocked in PBS containing 10 μg BSA/mL for 1 minute.

The inserts were then transferred to wells containing conditioned culture medium, i.e., medium collected from mTEC 3.10 control cell culture or mTEC 3.10 transfected with miR-155 mimic (cell cultures were grown for 24 hours). The conditioned medium was filtered through

0.45 μ filters before use. 100 µL of conditioned medium from the control cultures or the transfected cultures were placed in each well. A suspension containing 2.5 × 10^6^ thymocytes was then deposited on each of the inserts. The inserts were then placed inside the wells and incubated at 37°C in an atmosphere with 5% CO_2_ for 3 hours to allow the migration of thymocytes.

The inserts were then placed in wells containing 70% ethanol for 10 minutes to fix the thymocytes in passing through the pores. The outer surface of each insert was stained with Giemsa for 10 minutes and finally washed in deionized water to remove excess dye and then left to dry in room air. The inserts were visualized and photographed in a stereomicroscope model Stemi 508 Zeiss (Oberkochen, Germany).

The images were analyzed considering the average pixel of the migrated thymocytes (pixels/area). Thymocyte migration ranged from zero (no migration) to 255 pixels, which was the maximum pixel value for migrated and stained thymocytes. Migration values were compared between thymocytes that migrated in conditioned culture medium versus conditioned culture medium of cells transfected with miR-155 mimic.

## Ethics approval and consent to participate

This protocol was approved by the Ethics Committee on the Use of Animals in Research, Ribeirão Preto Medical School, certificate No. 003 / 2017-1.

## Consent for publication

Not applicable.

## Availability of data and materials

Numerical quantitative raw data from microarray hybridizations are available from the ArrayExpress repository (https://www.ebi.ac.uk/arrayexpress/) under accession number E-MTAB-7709.

## Competing interests

Authors declare no competing interests.

## Funding

This work was funded by Fundação de Amparo à Pesquisa do Estado de São Paulo (Fapesp, grant No. 17/1070-4 to GAP), Conselho Nacional de Desenvolvimento Científico e Tecnológico (CNPq, grant No. 305787/2017-9 to GAP), Coordenação de Aperfeiçoamento de Pessoal de Nível Superior (CAPES, financial code 001).

## Author’ s contributions

PPT: conceived the study, performed the experiments with mTECs including miR-155 mimic transfections, WB, microarray hybridizations and data analysis; EHO: conceived the study, performed the Aire-miRNA interaction prediction, cloned the Aire’ s mRNA 3′UTR, performed the luciferase reporter gene assay; MCV-M: performed the thymocyte migration assay; MJD and AFA: performed the microarray bioinformatics analysis; KFB-P: performed the immunofluorescence analysis of Aire protein; GAP: conceived the study, raised the hypothesis, designed the study workflow, provided the laboratory infrastructure and reagents, selected bioinformatics softwares, supervisioned the work and wrote the manuscript.

## Acknowledgments

We thank Dr W. Savino, Dr Daniela A. Mendes-da-Cruz and Dr Vinícius Cotta-Almeida (Fundação Instituto Oswaldo Cruz, Rio de Janeiro, RJ, Brazil) for their discussions and for giving us the mTEC 3.10 cell line. The HEK-293T cell line was kindly given by Dr. Fernando Q. Cunha (Ribeirão Preto Medical School, University of São Paulo, Brazil).

## Authors’ information

Not applicable.

---

**Supplemental Figure 1.**
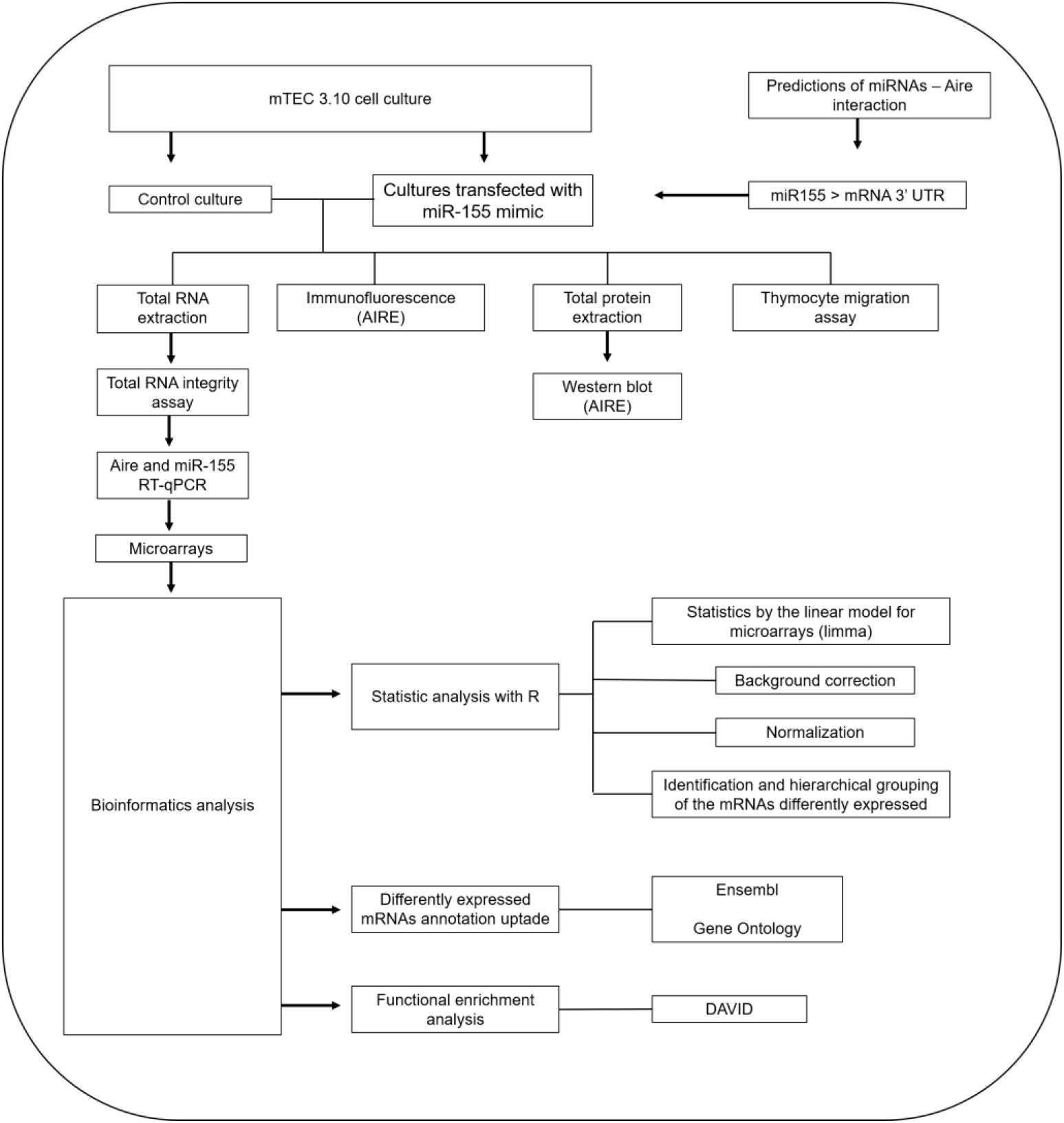
Flow diagram of the work.

## Notes

### Competing Interest Statement

The authors have declared no competing interest.

